# Water volume, biological and PCR replicates influence the characterization of deep-sea pelagic fish communities

**DOI:** 10.1101/2024.08.26.609755

**Authors:** Pedro A. Peres, Heather Bracken-Grissom

## Abstract

The pelagic deep sea is challenging to investigate due to logistical constraints regarding access and collection of samples, however environmental DNA (eDNA) can potentially revolutionize our understanding of this ecosystem. Although recent advancements are being made regarding eDNA technology and autonomous underwater vehicles, no investigation has been performed to assess the impact of different experimental designs using gear found on many research vessels (i.e., CTD mounted with Niskin bottles). Here, we investigated the effects of sampled water volume, biological and PCR replicates in characterizing deep-sea pelagic biodiversity at the level of species and exact sequence variants (ESVs, representing intraspecific variation). Samples were collected at 450m depth at night in the northern Gulf of Mexico using Niskin bottles, and we targeted the fish community using the MiFish primer (12S rRNA). Our results show that 1L is insufficient to characterize deep-sea pelagic fish communities. The 5L and 10L treatments detected similar community structure (i.e., the combination of number of species and relative occurrence) and numbers of species per biological replicate, but the 10L treatment detected a higher total number of species, more ESVs, and a different community structure when considering ESVs. We found that five biological replicates can detect up to 80% of the species detected in this study in the water collected in both 5L and 10L treatments. PCR replicates also had an important role in species and ESV detection, which implies increasing PCR replicates if water volume is limited. We suggest that future studies collect at least 5L, 5 or more field replicates, and 5-10 PCR replicates to adequately investigate deep-sea pelagic biodiversity using eDNA, considering resource limitations. Our study provides guidance for future eDNA studies and a potential route to expand eDNA studies at a global scale.

## INTRODUCTION

The use of DNA extracted from environmental samples (environmental DNA, eDNA) has become a common method to describe biodiversity and test ecological hypotheses (Dickie et al., 2018; Ruppert et al., 2019). Because this is a non-invasive approach, it has also gained popularity to inform conservation and management practices (Barnes & Turner, 2016; Beng & Corlett, 2020; Thomsen et al., 2024). eDNA can be particularly useful for documenting species diversity when a habitat is hard to access, or the target organisms are very small (Stat et al., 2017). Although this method is becoming a popular tool for coastal systems and empirical data has become more available, deep-sea studies are less numerous and, consequently, eDNA has been less employed to characterize deep-sea pelagic ecosystems (>200 m depth).

The deep-sea, including the midwater, is the largest habitat on Earth but one of the least understood (Ramirez-Llodra et al., 2010). Deep-pelagic animals participate in diel vertical migration, the largest migration on the planet in terms of biomass, and play a critical role in the biological pump and transport of nutrients, acting as the lifeline between the surface and deep ecosystems. In addition, there is potential economic importance represented by natural resources exploration, from fisheries to deep-sea mining (Thurber et al., 2014), although the impact of this exploitation needs to be carefully considered (Drazen et al., 2020; Haddock & Choy, 2024). Investigations of the deep sea and associated diversity are challenging due to logistical constraints regarding access, collecting, and filtering water samples. In this context, eDNA has the potential to revolutionize our understanding of this enigmatic and vast habitat.

Advances in robotics are improving sampling strategies, as seen using autonomous underwater vehicles (AUVs) in combination with *in situ* filtration for both surface and deep-pelagic communities (Billings et al., 2017; Yamahara et al., 2019; Yoerger et al., 2021; Govindarajan et al., 2022). However, this type of technology might not be accessible to many scientific groups due to the costs associated with specialized equipment, trained personnel, or because some models are custom-made and not commercially available, limiting the global use of this approach (Govindarajan et al., 2022). The limited working space of research vessels can also enhance these challenges if numerous materials and/or gear are needed for eDNA collection beyond what is available. We argue there is still a need to further investigate experimental designs that use common equipment with limited resources. This can facilitate the number of research groups performing eDNA in the deep sea, thus enabling large-scale collaborative studies.

Many eDNA studies, including the ones investigating the deep-pelagic habitat, use Niskin bottles coupled with a conductivity temperature depth (CTD) rosette to collect water samples. This piece of equipment is often provided on research vessels and coupled with other sensors to measure temperature, oxygen, salinity, and depth of the water column. Because the Niskin bottles can be triggered at a specific depth and location, researchers can design eDNA studies around the desired chemical, physical and /or biological properties of the ocean (McClenaghan et al., 2020; Govindarajan et al., 2021; Visser et al., 2021; Govindarajan et al., 2023). After collection, water samples are filtered, and the filters are preserved for future processing. Although there is some general guidance for eDNA studies (e.g., Dickie et al., 2018), little is discussed about specific sampling schemes focused on mid- and deep-water pelagic systems. If using Niskin bottles to collect water is a viable option, the amount of water and the number of biological replicates becomes a limitation because these factors are constrained by the size and number of bottles available. For instance, filtering many replicates of large amounts of water can be a laborious and sometimes impossible task (i.e. time constraints, filter clogging, and pump capacity). These limitations should be central in discussing deep-pelagic eDNA explorations as they will dictate the experimental design, representing a first and critical step in eDNA studies.

Besides equipment limitations, the pelagic deep-sea environment presents other unique challenges to consider when designing eDNA-based investigations. For instance, the magnitude of this habitat and often low density of individuals, can result in low levels of eDNA. Previous work has suggested that higher volumes of water, biological and PCR replicates might be necessary in offshore environments, where DNA can be sparsely distributed (Kumar et al., 2021; Cote et al., 2023); however, no experimental study has proposed or investigated potential guidelines (water volume, biological and PCR replication) for deep-pelagic biodiversity studies. Although obvious, one should not assume that sampling methods from coastal systems are sufficient or translatable for deep-pelagic waters. Such guidelines are essential for the development of eDNA investigations. In fact, efforts are being made towards creating standards in eDNA research at global and local scales (e.g., Environmental DNA Expeditions - UNESCO, unesco.org/en/edna-expeditions; National Strategy for Aquatic Environmental DNA – the White House Office of Science and Technology Policy, whitehouse.gov/wp-content/uploads/2024/06/NSTC_National-Aquatic-eDNA-Strategy.pdf).

Here, we investigate the current methodological knowledge gaps and provide guidance for future deep-pelagic eDNA studies. We test the impact of sampled water volume, biological and PCR replicates in assessing deep-sea fish pelagic biodiversity using equipment found on many research vessels (CTD rosettes and Niskin bottles). We collected 1L, 5L, and 10L simultaneously within the deep-scattering layer (450 m depth) in the northern Gulf of Mexico, which was targeted using acoustic methods. Each treatment consisted of biological replicates and PCR replicates per biological replicate, which were used to investigate species and exact sequence variant (ESV, representing intraspecific variation) detection, and community structure (i.e., a combination of species or ESV detection and relative occurrence). This experimental design allowed us to detect a preferred sampling scheme, acknowledging the challenges associated with pelagic deep-sea sampling and the limitations of equipment selection.

## MATERIALS AND METHODS

### Experimental design

We investigated the effects of water volume and biological and PCR replicates by examining samples collected simultaneously at 450m depth (see below). Our experimental design consisted of three treatments: 1L, 5L, 10L. Each treatment consisted of five biological replicates (n = 15) plus one negative control. The negative control is represented by 1L of ultrapure water filtered using the same apparatus that was used for the biological samples (see below). To investigate the effects of technical replicates, 10 PCR replicates were run for each biological replicate and the negative control when possible (n = 160). We had problems amplifying some 1L samples, resulting in few PCR replicates for two of the biological replicates. We decided to reallocate the resources by increasing one PCR replicate to the other biological replicates (Supplementary Table 1).

### Sampling and shipboard processing

Water samples were collected during a cruise on R/V Point Sur in the Northern Gulf of Mexico (29°00.175N, 87°31.131W) as part of the DEEPEND|RESTORE consortium (Cook et al., 2020). Samples were collected at 450m depth (nighttime) during a cast of a CTD (Sea-Bird 911plus) mounted with ten 10-L Niskin bottles. To maximize the detection of individuals, multifrequency acoustic backscatter data was collected with a calibrated pole-mounted echosounder system (Simrad EK60 and EK80). The transducers were mounted in an enclosed housing and suspended 2.5 m below the water surface (see Boswell et al., 2020 for additional details). Backscatter data were collected simultaneously at four frequencies (18, 38, 70, and 120 kHz). This was used to detect the deep scattering layer (DSL) in real-time and guide us in the precise moment to trigger the Niskin bottles. Because one of our objectives was to investigate the effect of filtering different water volumes, all samples were collected simultaneously during the same CTD cast to avoid any temporal or spatial variation in our results.

Once on board, seawater was transferred from Niskin bottles to 1-L sterile Nalgene bottles and immediately transported to the laboratory and filtered. We used a MilliporeSigma™ Chemical Duty Vacuum Pressure Pump (WP6111560) for filtration of all biological replicates and a negative control. Each replicate of specific seawater volume (1L, 5L, or 10L) was filtered through a sterile hydrophilic polyethersulfone (PES) filter (Pall Corporation; 47-mm diameter; 0.45-μm pore size). To prevent contamination, the filtering station was cleaned with 10% bleach, and nitrile gloves were worn throughout the filtering protocol and were changed frequently. Following sample filtration, the filters were placed in falcon tubes with silica beads, which function as a desiccator, drying out the filters and reducing DNA degradation. Samples were stored at −80°C in a freezer onboard. Upon arrival, samples were transported in dry ice to Florida International University and stored in −80°C freezer until sample processing.

### eDNA: Sample Processing, DNA extraction, PCR, sequencing

Genomic DNA from samples was extracted using the DNeasy Blood & Tissue Kit (250) (Cat. No. / ID: 69506) according to the manufacturer’s protocol. Whole filters (47 mm) were used for genomic DNA extraction. Genomic DNA was eluted into 200µl of AE buffer and frozen at - 20°C.

We decided to perform this investigation focusing on the fish community based on their abundance and ecological relevance in the deep-pelagic environment (Irigoien et al., 2014; Iglesias et al., 2023), and because of the availability of primers capable of giving species resolution in deep-sea fish communities (McClenaghan et al., 2020; Canals et al., 2021; Polanco F. et al., 2021). Portions of hyper-variable regions of the mitochondrial 12S ribosomal RNA (rRNA) gene were PCR amplified from each genomic DNA sample using the MiFishUF and MiFishUR primers with spacer regions (Miya et al., 2015). Both forward and reverse primers also contained a 5’ adaptor sequence to allow for subsequent indexing and Illumina sequencing. Each 25 µL PCR reaction was mixed according to the Promega PCR Master Mix specifications (Promega catalog # M5133, Madison, WI) which included 12.5µl Master Mix, 0.5 µM of each primer, 1.0 µl of gDNA, and 10.5 µl DNase/RNase-free H_2_O. DNA was PCR amplified using the following conditions: initial denaturation at 95°C for 3 minutes, followed by 45 cycles of 20 seconds at 98°C, 30 seconds at 60°C, and 30 seconds at 72°C, and a final elongation at 72°C for 10 minutes. PCR replicates were not pooled and were kept separate. Each reaction was visually inspected using a 2% agarose gel with 5µl of each sample as input to determine successful amplifications and amplicon size. Successful PCRs were cleaned by incubating amplicons with Exo1/SAP for 30 minutes at 37°C following by inactivation at 95°C for 5 minutes and stored at - 20°C.

A second round of PCR was performed to complete the sequencing library construct, appending with the final Illumina sequencing adapters and integrating a sample-specific,12-nucleotide index sequence. The indexing PCR included Promega Master mix, 0.5 µM of each primer and 2 µl of template DNA (cleaned amplicon from the first PCR reaction) and consisted of an initial denaturation of 95°C for 3 minutes followed by 8 cycles of 95°C for 30 sec, 55°C for 30 seconds and 72°C for 30 seconds. Final indexed amplicons from each sample were cleaned and normalized using SequalPrep Normalization Plates (Life Technologies, Carlsbad, CA). 25µl of PCR amplicon is purified and normalize using the Life Technologies SequalPrep Normalization kit (cat#A10510-01) according to the manufacturer’s protocol. Samples are then pooled together by adding 5µl of each normalized sample to the pool. Library preparation was performed by Jonah Ventures (Boulder, CO).

Sample library pools were sent for sequencing on an Illumina MiSeq (San Diego, CA) at the Texas A&M Agrilife Genomics and Bioinformatics Sequencing Core facility using the v2 500-cycle kit (cat# MS-102-2003) and a 15% PhiX spike in was added. Prior to sequencing, library size distribution was checked by fragment analysis prior to sequencing.

### eDNA: Bioinformatics

Raw sequence data were demultiplexed using pheniqs v2.1.0 (Galanti et al., 2021), enforcing strict matching of sample barcode indices (i.e, no errors). Cutadapt v3.4 (Martin, 2011) was then used to remove gene primers from the forward and reverse reads, discarding any read pairs where one or both primers (including a 6 bp, fully degenerate prefix) were not found at the expected location (5’) with an error rate < 0.15. Read pairs were then merged using vsearch v2.15.2 (Torbjørn et al., 2016), discarding resulting sequences with a length of < 130 bp, > 210 bp, or with a maximum expected error rate > 0.5 bp (Edgar & Flyvbjerg, 2015). For each sample, reads were denoised using the unoise3 denoising algorithm (Edgar, 2016) as implemented in vsearch, using an alpha value of 5 and discarding unique raw sequences observed less than 8 times. Counts of the resulting exact sequence variants (ESVs) were then compiled and putative chimeras were removed using the uchime3 algorithm, as implemented in vsearch. For each final ESV, a consensus taxonomy assignment was generated using a custom best-hits algorithm and a reference database consisting of publicly available sequences (GenBank, Benson et al., 2005) as well as Jonah Ventures voucher sequences records. The reference database was searched using the vsearch function usearch_global to conduct exhaustive pairwise semi-global alignments. Match quality was quantified using a custom, query-centric approach, where the % match ignores terminal gaps in the target sequence, but not the query sequence. The consensus taxonomy was then generated using the top hits for each ESV, defined as either all 100% matching reference sequences or all reference sequences with a % match greater than or equal to the highest % match minus 1. The consensus taxonomic assignment was then determined to be the highest taxonomic level for which there was at least 90% agreement among the top hits. Non-fish species were removed from the final dataset. For downstream analyses, we kept only ESVs that could be confidently assigned to a species (>98% similarity, Polanco F. et al.,2021). We kept as “sp.” species that had a reference deposited in public databases and an associated voucher number even if they were identified only to genus level (i.e., “genus name sp.”). The number of reads per species and ESV was coded as presence/absence. The final datasets consisted of 1) presence/absence of each species per biological replicate per treatment; 2) presence/absence of each species per PCR replicate per biological replicate per treatment; 3) presence/absence of each ESV per biological replicate per treatment; 4) presence/absence of each ESV per PCR replicate per biological replicate per treatment (Supplementary Table 1).

### Biodiversity analyses

We compared the number of species (richness) and ESVs detected per biological replicate in each treatment (1L, 5L, 10L) by performing a 1-way analysis of variance (ANOVA) followed by a posthoc Tukey’s HSD test. For comparisons of species and ESVs community structure across treatments, we performed a Permutation Multivariate Analysis of Variance (PERMANOVA; Anderson, 2001). First, we built a dissimilarity matrix based on the Bray-Curtis dissimilarities using the vegdist function in the vegan package (Oksanen et al., 2022). Because the package’s default is to use quantitative data for all indices, we set binary = TRUE to account for the presence/absence structure of our dataset. The resulting matrix was used as the response variable and volume as the explanatory variable in the PERMANOVA model using the adonis2 function. Significance was accessed using 9999 permutations. To investigate the effects of each treatment, we performed a pairwise permutational test for multiple comparisons set to 9999 permutations using the pairwise.adonis2 function in the pairwiseAdonis (Arbizu, 2020). The relationships among each biological replicate per treatment were visually represented by a Non-metric Multidimensional Scaling (nMDS) plot. This analysis was done using the metaMDS function implemented in the vegan package. Because convergence was not reached under default parameters, we used the argument trymax = 1000 to increase the number of random starts to find a stable solution. A permutation-based similarity percentage (PER-SIMPER) analysis was performed to identify the taxa that most contributed to the significant differences among treatments on both species and ESV datasets (Gilbert & Escarguel, 2019). We report the top 15 species indicated by the PER-SIMPER analysis, as this number corresponds to the number of species contributing to more than 33% (1/3) of treatment differences in our study. We also visualized the species and ESVs detected in each treatment by plotting a Venn Diagram using ggvenn package (Yan, 2023), and used the gplots package (Warnes et al., 2024) to find the exclusive and shared species/ESVs among treatments.

We investigated the efficiency of the number of biological replicates and PCR replicates in detecting species or ESVs by constructing rarefaction and extrapolation (R/E) plots using the package iNterpolation/EXTrapolation (iNEXT, Hsieh et al., 2016). We followed a non-asymptotic approach to plot a coverage-based R/E sampling completeness curve based on sampling-unit-based incidence data. Depending on the analysis, the sampling unit was considered the number of biological replicates (n = 5) per treatment (1L, 5L, 10L) or the number of PCR replicates (n = 10) per biological replicate (n = 5) per treatment (1L, 5L, 10L). R/E plots were constructed for both species and ESV datasets.

All analyses were run in R software (version 4.1.3).

## RESULTS

We retained 2,061,989 reads that were assigned to the Class Actinopteri. After all filtering steps, we retained 1,487,217 reads in total that could be assigned to a species (>98% similarity). The total number of reads (mean ± standard deviation) per treatment combining all biological replicates was 1L = 48,793.4 ± 49,311.5; 5L = 100,506.2 ± 27,211.7; 10L = 148,143.8 ± 51,240.5. The number of reads per biological replicate (mean ± standard deviation) per treatment was 1L = 4,900.3 ± 4,847.5; 5L = 9,136.9 ± 2,473.8; 10L = 13,467.6 ± 4,658.2. One biological replicate of the 1L treatment detected zero species. Negative controls had negligible amounts of contamination, with an average of 408.4 ± 1890.8 across PCR replicates. Only 6 species were found in the negative control, usually represented by one ESV in just one of the PCR replicates. We detected 98 taxa that could be confidently assigned to a species, representing a total of 661 ESVs (Supplementary Table 1). The total number of species (i.e., the sum across all replicates) detected by 1L = 39, 5L = 50, 10L = 79. The number of species detected per biological replicate per treatment (mean ± standard deviation) was 1L = 10.6 ± 12.34, 5L = 21.6 ± 6.87, 10L = 35 ± 8.74. The total number of ESVs detected by 1L = 112, 5L = 225. 10L = 426. The number of ESVs detected per biological replicate per treatment (mean ± standard deviation) was 1L = 35.6 ± 49.33, 5L = 84.6 ± 49.29, 10L = 182.4 ± 71.53.

The richness per biological replicate per treatment detected by the 1L treatment is lower than the number detected by the treatment filtering 10L (Tukey post-hoc, p = 0.004), but the number of species detected per biological replicate in 5L and 10L was not statistically different (p = 0.109; Table 1; Figure 2A). ESVs detected per biological replicate per treatment by 1L treatments are lower than the number of ESVs detected by the 10L treatment (p = 0.004); ESVs detected by 5L treatments are marginally lower than the 10L (p = 0.048; Table 1; Figure 2B). In both cases, 1L and 5L are not different (species, p = 0.206; ESVs, p = 0.399).

**Table 1.** Analysis of Variance (ANOVA) results comparing number of species (richness) per biological replicate found in each treatment. Permutation Multivariate Analysis of Variance (PERMANOVA) results comparing community structure among biological replicates per treatment (1L, 5L, 10L). DF: degrees of freedom; SS: sum of squares; F: F-value; Pseudo-F: pseudo-F-value; P: p-value.

| ANOVA - Richness |  |  |  |  |  |  |  |  |
| --- | --- | --- | --- | --- | --- | --- | --- | --- |
| Species |  |  |  |  | ESVs |  |  |  |
| Source of Variation | DF | SS | F | P | DF | SS | F | P |
| Volume | 2 | 1493 | 8.112 | 0.005 | 2 | 55860 | 8.394 | 0.005 |
| Residuals | 12 | 1104 |  |  | 12 | 3327 |  |  |

| PERMANOVA - Community Structure |  |  |  |  |  |  |  |  |
| --- | --- | --- | --- | --- | --- | --- | --- | --- |
| Species |  |  |  |  | ESVs |  |  |  |
| Source of Variation | DF | SS | Pseudo-F | P | DF | SS | Pseudo-F | P |
| Volume | 2 | 0.71959 | 1.8027 | 0.004 | 2 | 0.9926 | 1.3744 | 0.002 |
| Residuals | 12 | 2.395 |  |  | 12 | 5.3258 |  |  |

**Figure 1.**
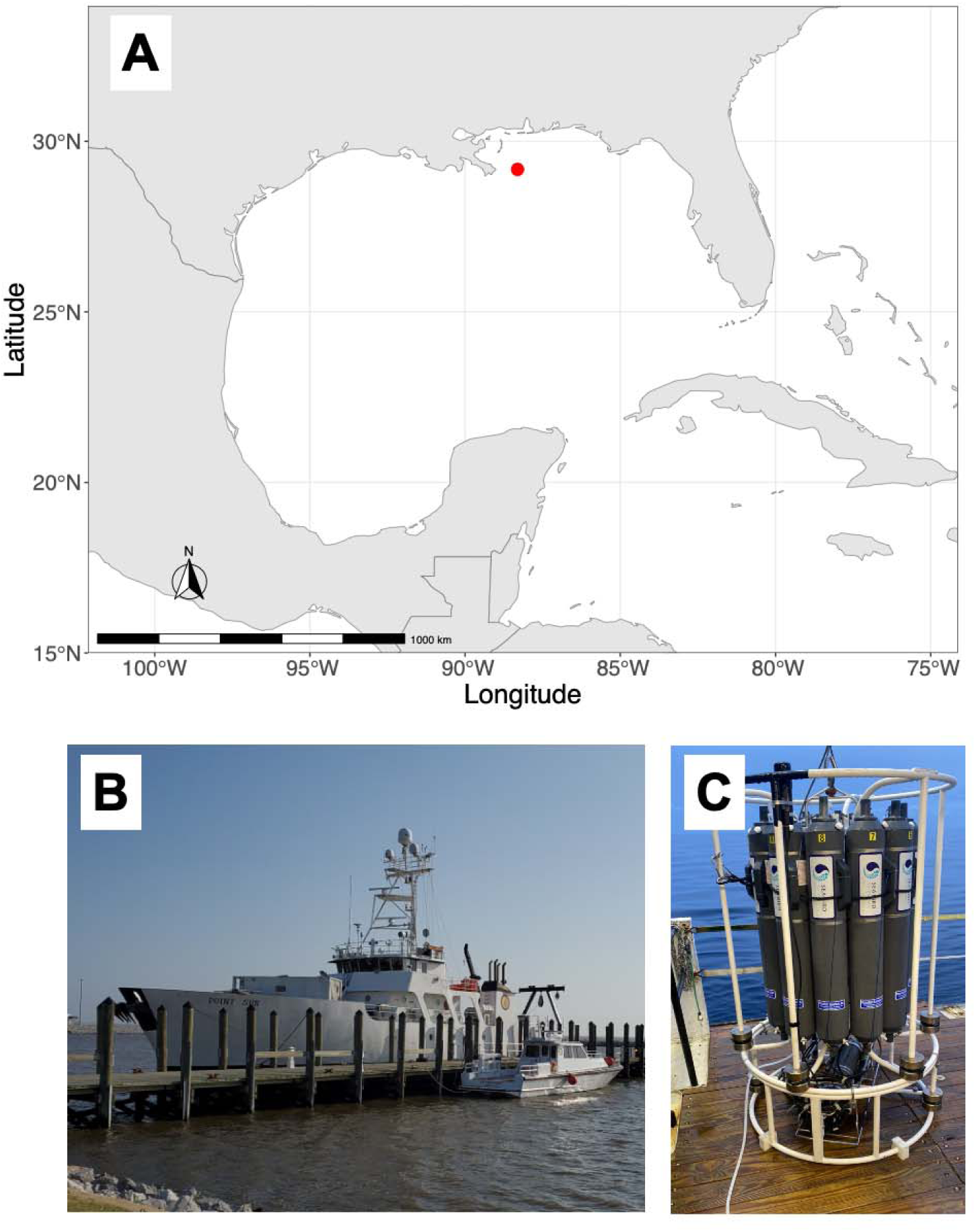
A) DEEPEND|RESTORE sampling site (DP08-01Aug22-B287N) in the northern Gulf of Mexico from where water was collected at 450m using a CTD. B) CTD and Niskin bottles used. C) R/V Point Sur.

**Figure 2.**
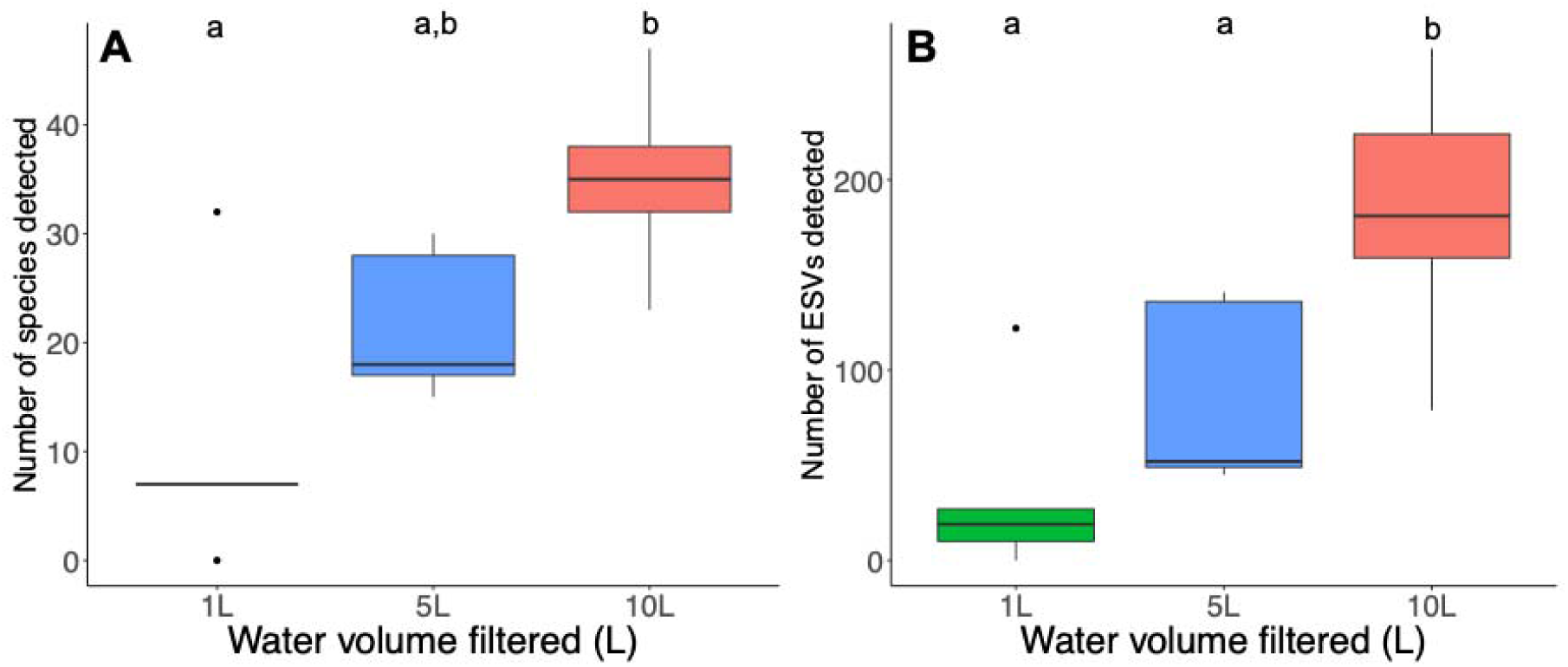
Box-plot representing the number of species (A) and ESVs (B) detected in each treatment (1L, 5L, 10L). Lower case letters on top of each box-plot represent the result of the *post-hoc* Tukey’s HSD test (similar letters are not statistically different). Colors represent different treatments. Green= 1L, Blue=5L, Red=10L.

Similarly, the PERMANOVA analysis indicated that volume affects community structure (Table 1). Considering the species dataset, the community detected by 1L marginally differs from the 5L (pairwise permutational test for multiple comparisons, p = 0.053) and differs from the 10L communities (p = 0.017), but 5L and 10L are not different (p = 0.113; 3A). Considering the ESVs dataset, each treatment detected a different community structure (1L vs. 5L, p = 0.04; 1L vs. 10L, p = 0.017; 5L vs. 10L, p = 0.031, 3B). When looking at the species contributing to the significant differences found in each comparison, the PER-SIMPER analysis indicates the top 15 species contributed 46.8% to the dissimilarity between 1L vs. 5L, and 34.2% between 1L vs. 10L (Table 2), being all of them more likely to be found in the 5L and 10L than in the 1L treatment. Although each species contributes little to the dissimilarity found, PER-SIMPER indicates that 8 out of the 15 species had a role in differentiating 1L from 5L and from 10L: *Barbourisia rufa, Benthosema suborbitale, Caranx crysos, Cyclothone pallida, Diaphus lucidus, Gonostoma elongatum, Lampanyctus alatus,* and *Rondeletia loricata.* Twenty-two species (22.4%) were found in all treatments (Figure 4, Supplementary Table 1). Thirty-two species were exclusively detected in the 10L treatment (Supplementary Table 1), with twenty-two of them (68%) detected in only one replicate. In the other two treatments (1L and 5L), all exclusive species are represented by detections in only one biological replicate (1L = 11, 5L = 7, Supplementary Table 1). Regarding ESVs, the top 15 ESVs contribute to 33.6% (1L vs. 5L), 30.7% (1L vs. 10L), 25.6% (5L vs. 10L) (Supplementary Table 1). Most of the ESVs contributing to the differences between 1L vs. 5L and 10L indicated by PER-SIMPER are ESVs from the species detected in the species-level analysis (Table 2, Supplementary Table 1). In the case of the 5L vs. 10L comparison, all ESVs represent species shared between the two treatments (Supplementary Table 1). Only twenty-seven ESVs were shared among all treatments (4.1%), and the 10L treatment had the highest number of exclusive ESVs (351, 53.3%) (Figure 4B)

**Table 2.** Permutation-based similarity percentage (PER-SIMPER) results showing the top 15 species contributing to the differences found between statistically different communities (see PERMANOVA results).

| 1L vs. 5L |  | 1 vs. 10L |  |
| --- | --- | --- | --- |
| Species | % | Species | % |
| *Cyclothone pallida* | 4.3 | *Lagocephalus laevigatus* | 3.2 |
| *Gonostoma elongatum* | 4.3 | *Canthigaster rostrata* | 2.8 |
| *Diaphus lucidus* | 3.6 | *Cyclothone pallida* | 2.8 |
| *Caranx crysos* | 3.3 | *Diaphus lucidus* | 2.8 |
| *Cyclothone pseudopallida* | 3.3 | *Gonostoma elongatum* | 2.8 |
| *Barbourisia rufa* | 3.3 | *Benthosema suborbitale* | 2.3 |
| *Lampanyctus alatus* | 3.1 | *Notolychnus valdiviae* | 2.3 |
| *Melamphaes simus* | 3.1 | *Caranx crysos* | 2.1 |
| *Diaphus perspicillatus* | 3.0 | *Lampanyctus alatus* | 2.1 |
| *Cyclothone obscura* | 2.8 | *Barbourisia rufa* | 2.0 |
| *Rondeletia loricata* | 2.8 | *Cyclothone acclinidens* | 1.9 |
| *Benthosema suborbitale* | 2.5 | *Rondeletia loricata* | 1.9 |
| *Lepidophanes guentheri* | 2.5 | *Nannobranchium lineatum* | 1.9 |
| *Seriola rivoliana* | 2.4 | *Caranx hippos* | 1.8 |
| *Diaphus rafinesquii* | 2.4 | *Lutjanus campechanus* | 1.8 |
| Total | 46.8 |  | 34.3 |

**Figure 3.**
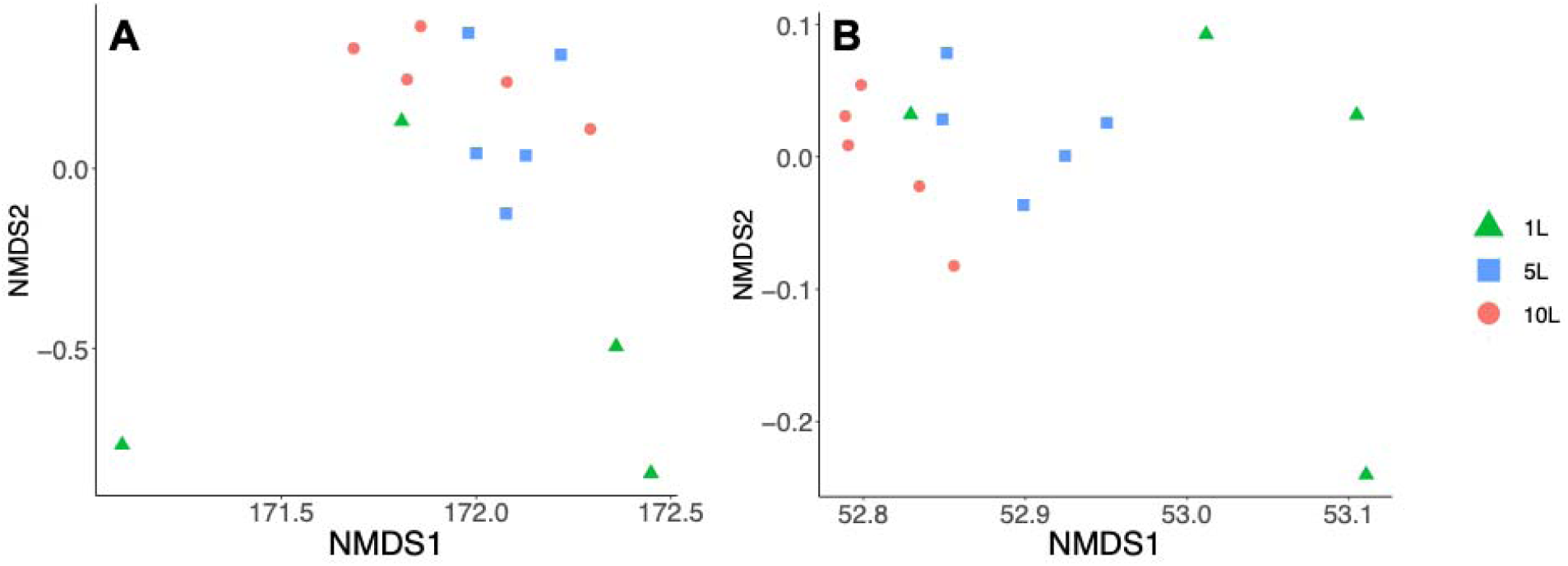
Non-metric multi-dimensional scaling (NMDS) plot representing the species (A) and ESVs (B) community structure found in each treatment (1L, 5L, 10L). The 1L replicate that detected zero species/ESVs was excluded to facilitate visualization of the spatial distribution of samples. Colors represent different treatments. Stress: A = 7.716376e-05; B = 8.470151e-05.

**Figure 4.**
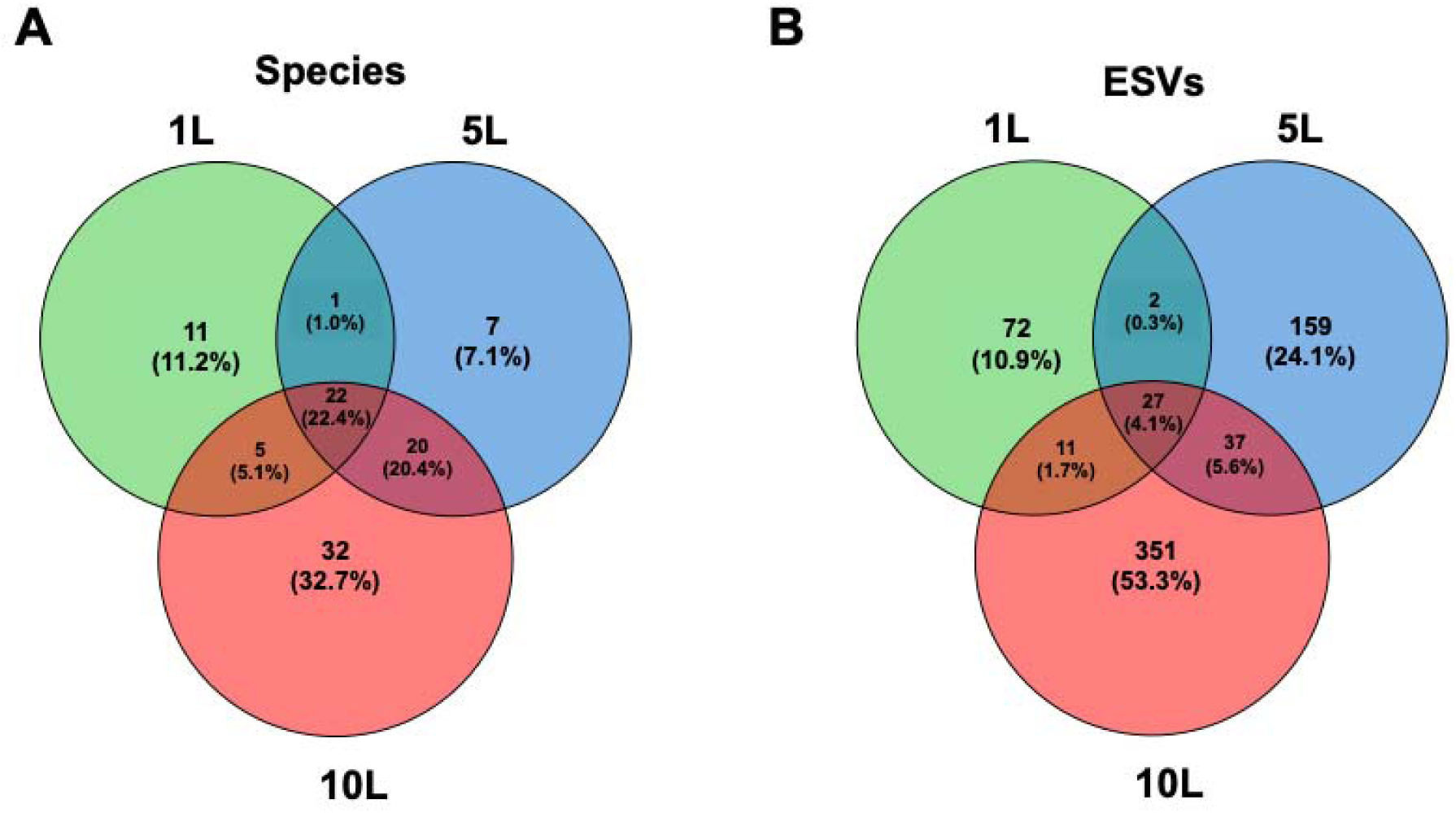
Venn diagram indicating the number of exclusive species in each treatment (1L, 5L, 10L), and shared species between two treatments or shared among all treatments (A); Venn diagram indicating the number of exclusive ESVs in each treatment (1L, 5L, 10L), and shared ESVs between two treatments or shared among all treatments.

When analyzing the impact of the number of biological replicates in detecting the total biodiversity across all of our samples (i.e., all the species detected in this study), we show that 5 biological replicates could detect up to 80% of all 98 species in both 5L and 10L treatments. The extrapolation analysis indicates that 10 replicates would recover nearly 100% of the species detected in the water collected (note: not total biodiversity from the region, but all 98 species). In contrast, 5 biological replicates of 1L treatment detected only 50% of the total biodiversity, and the extrapolation analysis indicates that more than 10 biological replicates would be necessary to detect 100% (Figure 5A). A different scenario is found when analyzing ESVs, showing that 5 biological replicates detected ∼40% of the 611 ESVs detected. Extrapolation analyses indicated that more than 10 biological replicates would be necessary to detect all ESVs (Figure 5B).

**Figure 5.**
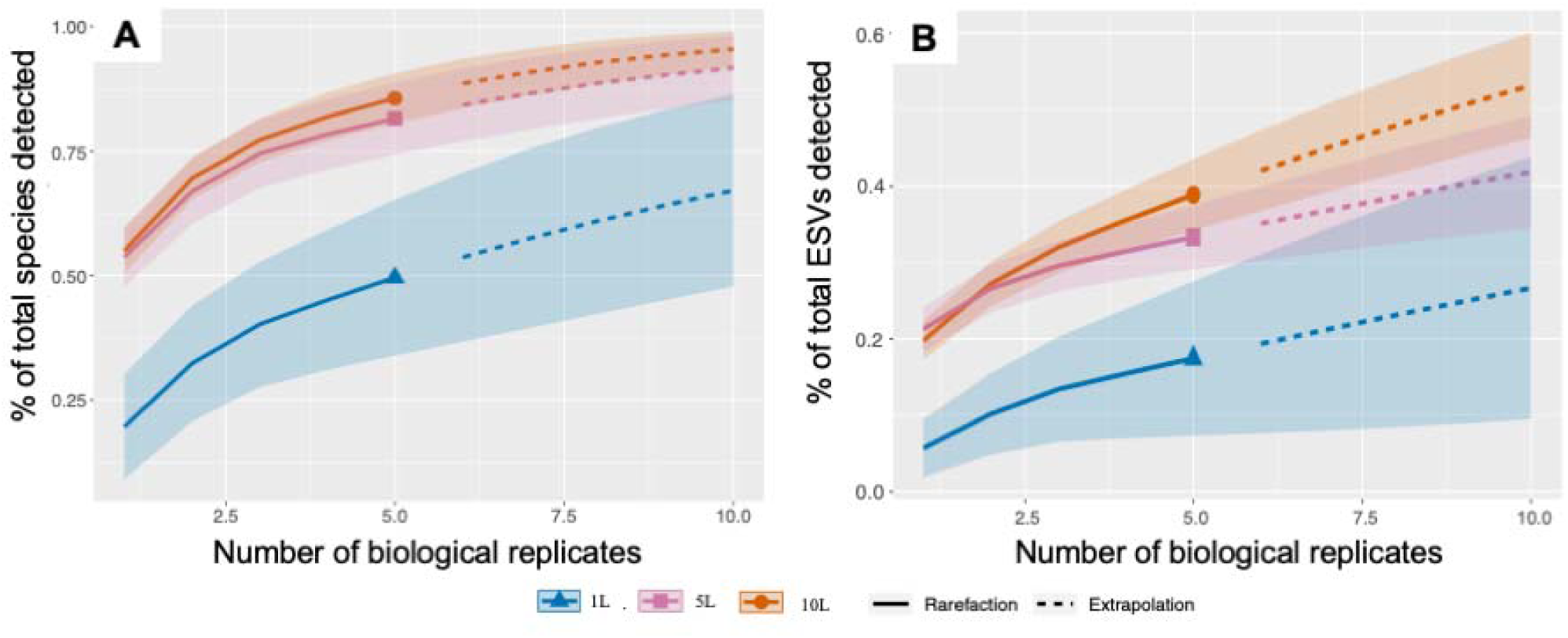
Rarefaction and extrapolation plot depicting the increase of species (A) or ESVs (B) detected as more biological replicates were added per treatment (1L, 5L, 10L). Shaded areas represent standard error.

When analyzing the impact of the number of PCR replicates per biological replicate, we find that 10 PCR replicates show variable biodiversity coverage for 5L, detecting between 60-85% of total number of species depending on the biological replicate (Figure 6B). Similarly, 10L biological replicates show a similar pattern, but detecting around 80-90% of the total number of species (Figure 6C). In contrast, results concerning PCR replicates of 1L treatments are extremely variable (Figure 6A). Similar to the previous results, PCR replicates R/E plots analyzing ESVs indicate large variation among 1L biological replicates (Figure 6D), and moderate/low variation among 5L and 10L biological replicates. In these cases, 10 PCR replicates detected between 20-60% of the ESVs sampled in this study (Figure 6E-F). One of the biological replicates in the 1L treatment did not have enough DNA for running all PCR replicates.

**Figure 6.**
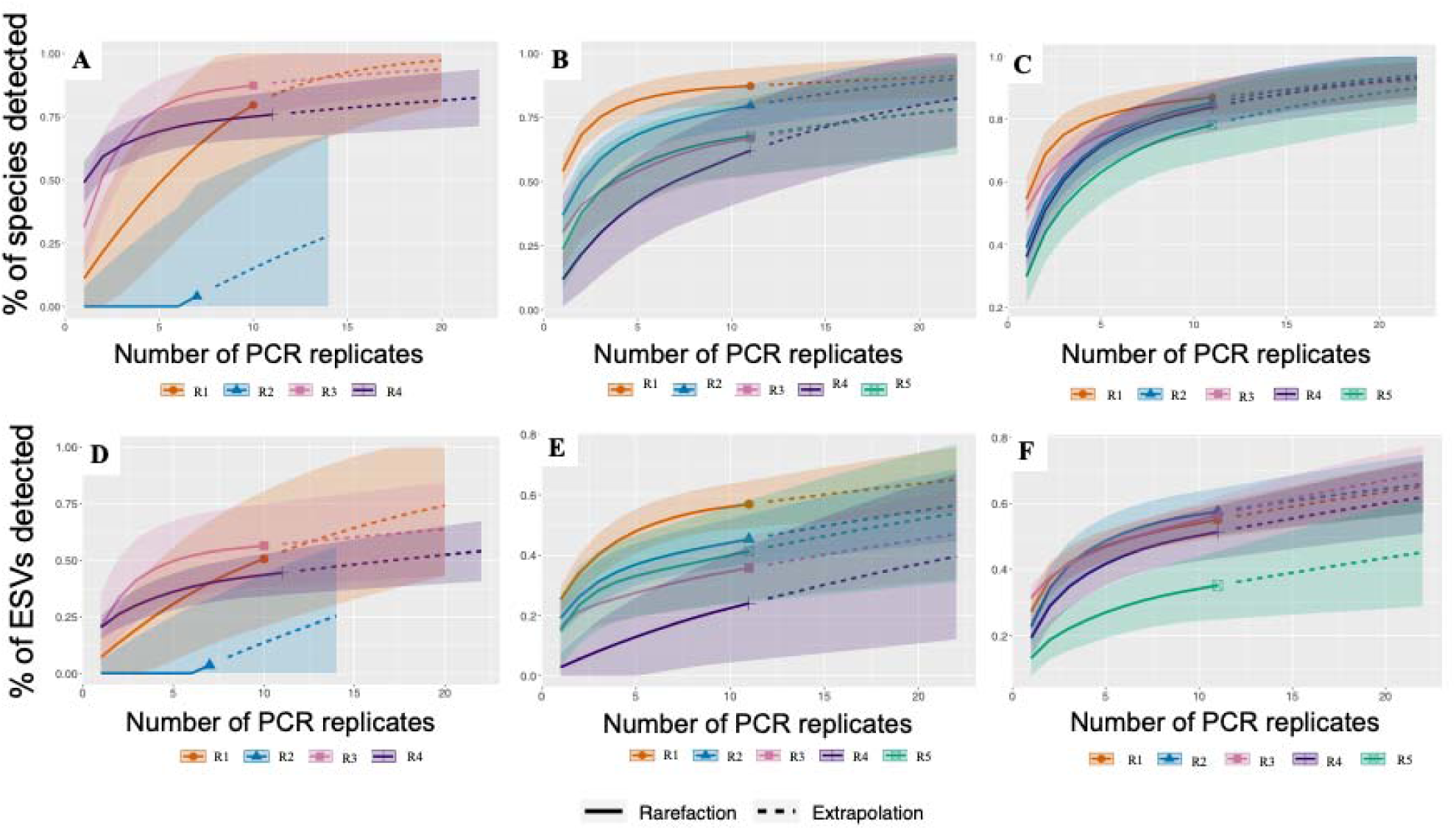
Rarefaction and extrapolation plot depicting the increase of species/ESVs detected as more PCR replicates were added biological replicate per treatment (A: species-1L; B: species-5L; C: species10L; D: ESVs-1L; E: ESVs-5L; F: ESVs-10L). Shaded areas represent standard error.

## DISCUSSION

Experimental design is a crucial part of any ecological study, but there are no golden rules when implementing eDNA approaches (Deiner et al., 2017; Dickie et al., 2018; Ruppert et al., 2019; Cote et al., 2023). Sampling design considerations must be carefully considered before going to sea, especially for deep-pelagic eDNA investigations. Our study shows that when using Niskin bottles coupled with CTD, filtering 5L and 10L did not result in statistically different species community structure (i.e., species detection combined with relative occurrence) but detected different ESV community structure. However, the total number of species and ESVs (i.e., the sum of all species/ESVs detected across replicates) is higher in the 10L treatment. The number of biological and PCR replicates impacted species and ESV detection, which we will expand on below. As far as we know, previous investigations have not tested how different sampling schemes (water volume and replication) using common equipment (e.g., Niskin bottles coupled with CTD) can impact deep-sea biodiversity assessments. Our results provide empirical evidence on how different experimental designs can affect deep-pelagic eDNA investigations using Niskin bottles coupled with CTD. Ultimately, our work can help future eDNA studies make informed decisions considering the pros and cons of alternative sampling strategies, potentially aiding in expanding the number of comparable datasets.

### Water volume

In aquatic eDNA studies, a standing question is: how much water should be filtered? Data from studies conducted in shallow systems indicate that filtered water volume is known to impact taxa detection (Muha et al., 2019; Kumar et al., 2022). In these cases, filtered water volume usually varies from 0.5 mL to 2L (Rees et al., 2014; Bessey et al., 2020; Kumar et al., 2022). Few studies have investigated the impacts of different filtering protocols, resulting in similar water volumes being filtered between shallow water and deep-pelagic eDNA studies (McClenaghan et al., 2020; Diao et al., 2023). Directly transposing shallow water standards to mid- and deep-pelagic ecosystems eDNA investigations might not be ideal considering the differences between environments. Although none of the treatments alone could detect all the 98 species and 661 ESVs found in this study, our results indicate that 1L is far from ideal for characterizing deep-pelagic fish communities. In terms of total number of species detected, the 1L treatment detected 22% and 51% less species than the 5L and 10L, respectively. Regarding the total number of ESVs, the 1L treatment detected 51% and 74% fewer ESVs than the 5L and 10L, respectively. Therefore, filtering 1L of water seems to fail in detecting many species and most of the within species variation (because species could be represented by multiple ESVs, indicating intraspecific diversity). This can be explained by the low eDNA abundance per volume of water in deep-pelagic habitats, leading to stochastic detection of eDNA molecules in 1L samples. The open ocean is the largest environment on Earth, covering 70% of our planet, and the deep sea represents 80-90% of our oceans, making it the largest habitat on Earth (Ramirez-Llodra et al., 2010). Therefore, species abundance and distribution are expected to differ substantially across spatial scales when comparing shallow- and deep-pelagic ecosystems.

When comparing community structure (i.e., species detection combined with relative occurrence), 5L and 10L treatments are not statistically different, indicating that treatments shared many species and differences in total number of species are represented by species with low detection rates. Community structure differences between 1L vs. 5L and 10L can be explained by the omission of rare species, but also species known to be very common in deep-pelagic habitats within the Gulf of Mexico based, including species from Gonostomatidae (*Cyclothone pallida*, *Gonostoma elongatum*) and Myctophidae (*Lampanyctus alatus*) (Ross et al., 2010)). These species are commonly collected as part of paired net sampling (eDNA and nets) using the MOCNESS 10 in the same sampling localities (personal observation – unpublished data). It is important to notice that we are discussing these data under a community structure framework, with the goal of comparing samples across time and space looking at species detection and how frequently each of them is detected. When the main goal of an investigation is to detect one or a few specific and/or rare taxa, or maximize species detection aiming the creation of species lists, higher volumes should be aimed as DNA detection likely increases (Sepulveda et al., 2019; Bessey et al., 2020; Govindarajan et al., 2022). As mentioned, our study also reports evidence for this latter statement as the 10L detected a total number of species higher than 1L and 5L treatments (but the number of species per biological replicate is not different from the 5L treatment, see next section).

A comparison of the number of operational taxonomic units (OTUs) found in different filtered water volumes (∼5-60L) also indicated no changes in total OTUs, although this varied according to location (Yoshida et al., 2023). A similar conclusion was reported when comparing CTD samples (∼2L) and a recently developed large-volume autonomous eDNA sampler with in-situ filtration mounted on the midwater robot Mesobot (∼40-60L): the community structure detected was not different between CTD and the Mesobot samples (Govindarajan et al., 2022). Species detection might be driven by their relative abundance in the sample, meaning that more water filtered does not necessarily affect the community structure, even if more rare species are detected. These rare species have a minor role when discussing communities’ similarities/differences. We should notice that this is related to the eDNA signal detected, but some species might not be rare in terms of abundance at the organismal level. Species detection based on eDNA is constrained by the persistence of DNA molecules in the water, correct amplification via PCR, and reference libraries that contain the necessary information to identify species correctly.

Our study also investigated intraspecific variation by looking at ESVs, representing within species genetic variation. Our findings indicate that the 10L treatment detected more ESVs and a different ESV community structure than the 1L and 5L treatments. This is similar to Govindarajan et al. (2022), who showed that the total number of sequence reads and ESVs was higher in samples collected by an autonomous sampler capable of filtering higher volumes of water (Govindarajan et al., 2022). Higher volumes also detected more ESVs in a study comparing 250mL and 1.5L collected using a CTD (McClenaghan et al., 2020). Therefore, choosing how many liters of water will be filtered should also consider whether the detection of more ESVs is part of the goal of the study. For instance, recent analytical developments have suggested that the number of intraspecific ESVs can be used to estimate species abundance (Yoshitake et al., 2019).

The understanding of the impacts of filtered volumes on the characterization of deep-pelagic communities is still under development, and more studies will illuminate whether there is an ideal volume that can be filtered. We should also note that Govindarajan et al. (2022) targeted the 18S rRNA region and many metazoan groups at lower taxonomic resolution, while we targeted 12S rRNA and fish species at a higher taxonomic resolution. The taxonomic resolution and efficiency vary according to the primer used and the region targeted (Ruppert et al., 2019; Zhang et al., 2020; Polanco et al., 2021; Kumar et al., 2022), being an important step during experimental design. We used a taxon-specific primer (MiFish) to better understand the nuances of sampling schemes at the highest taxonomic resolution (i.e., species). It is not possible to say if the results of Govindarajan et al. (2022) would be different whether they had targeted a different region (i.e., used a different primer) or investigated higher taxonomic resolution.

Our results are relevant when considering the effort-time-cost balance related to eDNA fieldwork that does not involve an autonomous water sampler and specialized filtering apparatus. Collecting and filtering water can be laborious and time-consuming and involve multiple researchers when using commonly available gear, like Niskin bottles and vacuum pumps. Here, we suggest that, when limitations to water collection exist, efforts toward increasing filtered water volume above 10L may not impact deep-pelagic community comparisons under a community structure framework, although more species can be detected. Therefore, alternative sampling strategies should be considered in light of the questions to be answered in deep-sea eDNA investigations.

### Biological and PCR replicates

Another crucial factor to consider when performing eDNA investigations is the number of biological and PCR replicates. Our work is one of the first to address the variation in these factors when characterizing deep-sea systems. Although some comparisons are not statistically significant, NMDS plots and rarefaction curves suggest large variation among 1L biological replicates in comparison to 5L and 10L treatments. Five biological and 10 PCR replicates of the 1L treatment were insufficient to assess the local fish biodiversity (i.e., the species and ESVs detected in this study), and even our extrapolation analyses could not reach levels compared to the other treatments. This means that not only the volume of water but also the number of replicates is important when considering eDNA applied to mid- and deep-water pelagic systems.

A striking conclusion is that 10L did not statistically detect more species per biological replicate than 5L, even though the mean number of reads in the 10L treatment was almost 50% larger than in the 5L treatment. However, the 10L treatment statistically detected more ESVs per biological replicate (i.e., more ESVs per species, representing intraspecific variation), and higher total number of species (i.e., the sum of all species detected across replicates). We should acknowledge the lack of differences in the number of species detected per biological replicate could have been impacted by our sample size. In fact, although not statistically different, Figure 2A shows a likely positive trend in higher number of species detection per biological replicate following increased water volume filtration, so results should be interpreted with caution. An optimal deep-sea eDNA experimental design could aim for a minimum of five biological replicates of a minimum of 5L, which is doable considering most traditional water samplers (e.g., McClenaghan et al., 2020; Feng et al., 2022; Govindarajan et al., 2023). Our results indicate that, under certain circumstances, it could be adequate to favor less volume and more biological replicates if one is limited on the amount of water that they can sample using their available gear (e.g., Niskin bottles coupled with CTD can only sample a specific total volume of water). Future studies should address the trade-offs between water filtered and the number of biological replicates using autonomous samplers.

Similar to the optimal number of biological replicates, the investigation of the ideal number of PCR replicates in eDNA studies is still scarce in the literature. We provide empirical data that reinforce previous suggestions that recommend increasing PCR replicates to improve detection probabilities and fish biodiversity estimates (Bessey et al., 2020; Stauffer et al., 2021; Cote et al., 2023; Yoshida et al., 2023). Amplification is a stochastic process during PCR reactions, and more reactions will probably result in the detection of more taxa by chance. The exact number needed will vary depending on the probability of DNA detection, which varies according to species abundance, rate of shedding DNA, environmental conditions, and other factors (Ruppert et al., 2019). A simulation study showed that four PCR replicates might result in high levels of false negatives, and six to eight PCR replicates should be performed depending on the probability of DNA detection (Ficetola et al., 2015). As discussed before, it is feasible to assume that DNA is sparsely distributed in the deep sea, considering this is the largest habitat on Earth and individuals’ density might not be high. Still, PCR replicates are often low in studies investigating deep waters (2-4 PCR replicates; e.g., McClenaghan et al., 2020; Canals et al., 2021; Govindarajan et al., 2022; Cote et al. 2023; Yoshida et al., 2023). Our results show that all samples benefited from more PCR replicates when considering species and ESVs detection. Besides biological replicates, the ideal sampling scheme should balance the volume of water filtered with the number of PCR replicates. That is, if limited by the water volume, the best strategy is to increase the number of PCR replicates; if not limited by water volume, fewer PCR replicates can be performed. Ideally, future studies should first find the optimal design to maximize water volume and PCR replicates accordingly.

We should acknowledge that increasing the number of replicates and PCR replicates can rapidly increase the cost of a project. Acquiring an up-to-date technological eDNA sampler can be expensive, but it can come at the cost of potentially having fewer biological and PCR replicates. Future studies should explore if fewer replicates of high-volume samples have the statistical power to test ecological hypotheses. Based on our results, a sampling strategy of 5L, 5 biological replicates, 5 PCR replicates can detect between 45-80% of the total number of species/18-50% of the total number of ESVs per biological replicate using Niskin bottles and CTD. Increasing any of the factors can be beneficial and is recommended if possible, but following this guideline represents a balance between water volume filtered, project cost, feasible characterization of the community under investigation, and an adequate number of samples to have enough statistical power to test ecological hypotheses.

### Limitations

It is important to mention the limitations of equipment selection in deep-sea eDNA studies. For instance, for CTDs coupled with Niskin bottles, the maximum water volume is limited by the capacity and number of bottles attached to the CTD, the number of deployments within a scientific mission, and cross-contamination from different casts. Our study leverages gear often found on research vessels and could not test if filtering more than 10L would change the biodiversity characterization, especially at the species level, as ESV diversity is favored by filtering more water. A potential future standard in this type of study could be first to investigate the ideal water volume and replicates needed to characterize the community at the specific ocean basin using a specific primer. Considering our results and Govindarajan et al. (2022), 5L-10L might be sufficient for the northern Gulf of Mexico when targeting the 18S and 12S rRNA markers. We should note that some taxa might shed less DNA than others (Andruszkiewicz Allan et al., 2021), which should also be considered when planning an eDNA investigation. For instance, we could not detect substantial decapod eDNA using the MiDecap primer set (Komai et al., 2019) using the volume of water and replicates used in this study (data and results not shown). This is surprising, considering the diversity and abundance of pelagic decapods in the Gulf of Mexico collected using nets and with available reference sequences (Burdett et al., 2017; Varela et al., 2021). We predict that primer compatibility and shedding may have been involved in the lack of recovery, but further studies should explore the impact of water volume and reference libraries on the recovery of deep-sea crustaceans and other taxa. Simulation analysis indicates that increasing the water volume filtered could lead to similar species being detected using eDNA and net sampling, reinforcing this idea (Cote et al., 2023). Additionally, if detecting specific taxa or species (instead of community structure) is the goal, one might consider maximizing the water volume collected, as mentioned before, or employing qPCR instead of metabarcoding approaches (Sepulveda et al., 2019; Bessey et al., 2020; Govindarajan et al., 2022). Still, the exact volume might be taxa-specific and should also be a first step in this type of investigation. Another factor to be considered is that eDNA might degrade while samples are waiting to be or while they are being processed (Holman et al., 2022). In our case, all replicates were filtered in a randomized order to avoid this pitfall. However, the decay rate of deep-sea eDNA samples is still unknown for many taxa, and future studies should address this knowledge gap (but see McCartin et al., 2022 for insights).

Besides the technical limitations mentioned, the primary pitfall of all deep-water eDNA investigations is the catastrophic lack of reliable reference libraries. eDNA approaches are 100% reliant on the completeness and quality of available reference databases for properly identifying taxa (Bylemans et al., 2018; Collins et al., 2019; Schenekar et al., 2020). In our study, ∼50% of our reads could not be confidently assigned to a fish species. Another situation that highlights the importance of reliable reference libraries is that we also detected a few Indo-Pacific species in our final dataset. We cannot confirm if they represent real occurrences or problems in reference sequences available in public databases. This study, among others, highlights the urgent need for deep-pelagic reference libraries that are linked to curated, voucher specimens deposited in museums that can be verified by taxonomic experts. Current efforts, including our own, are expanding deep-sea reference libraries (e.g., Bucklin et al., 2010; Weigand et al., 2019; Varela et al., 2021; Govindarajan et al., 2022), which need to be standardized, shared and extended to as many taxa and locations as possible.

### Conclusions

The application of eDNA to mid- and deep-water ecosystems is gaining popularity, however few studies have experimentally tested the effects of different sampling protocols on the recovery of species biodiversity (Canals et al., 2021; Easson et al., 2020; Govindarajan et al., 2021; Laroche et al., 2020; Merten et al., 2021). Our study provides empirical results on how experimental design can impact the characterization of deep-pelagic communities. Specifically, we indicate that a minimum of 5L-10L of water should be filtered when targeting the deep-scattering layer and paired with a minimum of 5 or more biological replicates for deep-sea eDNA studies when investigating community structure. Additionally, we show the importance of more PCR replicates when analyzing deep-sea samples. Technological advances in eDNA samplers are crucial and are expanding our understanding of deep-sea ecosystems. But, until eDNA samplers capable of filtering higher volumes of water with *in-situ* preservations become accessible for all, using Niskin bottles coupled with a CTD can be a way to facilitate midwater eDNA investigations. Most importantly, because deep-ocean explorations are constrained by logistics and budget, we believe our work can help in providing some guidance to increase local and global-scale observations.

